# Synaptojanin1 deficiency upregulates basal level autophagosome formation in astrocytes

**DOI:** 10.1101/2021.01.08.425969

**Authors:** Ping-Yue Pan, Justin Zhu, Asma Rizvi, Xinyu Zhu, Hikari Tanaka, Cheryl F. Dreyfus

## Abstract

Macroautophagy (hereafter, autophagy) dysregulation is implicated in multiple neurological disorders. While the autophagy pathways are heavily investigated in heterologous cells and neurons, how autophagy is regulated in the astrocyte, the most abundant cell type in the mammalian brain, is less understood. Here we report that Synaptojanin1 (Synj1), a neuron enriched lipid phosphatase, is expressed in low levels in astrocytes and represses autophagy at the basal level. Synj1 is encoded by the *Synj1* gene, whose missense mutations are linked to Parkinsonism with seizure. While the best-known role of Synj1 is to facilitate synaptic vesicle recycling, recent studies suggest that Synj1 also regulates autophagy. Our previous study using the *Synj1* haploinsufficient (*Synj1+/−*) mouse demonstrated that *Synj1* deficiency was associated with an age-dependent autophagy impairment in multiple brain regions. We now use cultured astrocytes from *Synj1* deficient mice to investigate its role in astrocyte autophagy. We demonstrate that Synj1 deficient astrocytes exhibit increased LC3 puncta, which is more pronounced when lysosomal acidification is blocked. The increased autophagosome formation is accompanied by reduced autophagy substrate, p62, but an insensitivity to starvation induced autophagy clearance. Moreover, we show, for the first time, that the Parkinsonism associated R839C mutation impacts astrocyte autophagy. The profound impact of this mutation on Synj1’s phosphatase functions results in elevated basal level autophagosome formation and clearance that mimics *Synj1* deletion. We find that energy sensing molecules, including mTOR and AMPK, are altered in *Synj1* deficient astrocytes, which may contribute to the enhanced basal level autophagy.

## Introduction

Regulated membrane trafficking is essential for the function of neuron and glia. Autophagy is part of the intricate membrane trafficking network and is often known as the self-eating process that maintains cellular homeostasis and copes with energy crisis. The formation of the autophagosome, a double-membraned structure, is typically induced by nutrient deprivation/starvation, and the sequestered cellular components (damaged organelles or protein aggregates) are eventually degraded in the autolysosome. Autophagy dysregulation has been implicated in various neurodegenerative disorders [1, 2], and the impairment in autophagy clearance is thought to contribute significantly to the accumulation of various forms of protein aggregates found in Alzheimer’s disease, Parkinson’s disease and Huntington’s disease. Emergent evidence suggests that there is a presence of protein aggregates in astrocytes as in neurons [3, 4], which highlights the importance of astrocyte autophagy in disease progression [5, 6]. While autophagy regulation has been studied mostly in heterologous cells and neurons, the process is not well understood in astrocytes, and its distinctive regulatory mechanisms have only begun to be recognized.

Synaptojanin1 (Synj1) is one of the key proteins involved in cellular trafficking. For the past two decades, the best-known function of Synj1 has been to facilitate neuronal synaptic vesicle (SV) recycling primarily through regulating the conversion of membrane phosphoinositide [7–10]. Synj1 contains two highly conserved inositol phosphatase domains: the SAC1-like domain hydrolyzes PI4P [11] as well as the 3’ phosphate on PI3P and PI(3,5)P_2_ [12], while the 5’-phosphophatase domain is a more potent enzyme that hydrolyzes the 5’ phosphate on the PI(4,5)P_2_ and PI(3,4,5)P_3_. The proline-rich domain (PRD) of synj1 is more variable and is subject to active phosphorylation and protein interaction [13–16]. Variation in the PRD results in two Synj1 isoforms [17]. The 170 kDa long isoform is ubiquitously expressed, whereas the 145 kDa short isoform is known to be enriched in neurons and particularly at the presynaptic terminals for SV recycling.

Deletion of *Synj1* results in an accumulation of clathrin coated pits at the presynaptic terminal and produces a lethal phenotype at birth [9, 10]. In the recent decade, multiple neurological disorders have been linked to the dysregulation of the *SYNJ1* gene. For example, *SYNJ1* gene triplication or overexpression leads to early endosome enlargement, hippocampal dysfunction and cognitive impairments, which may contribute to early onset Alzheimer’s disease and Down syndrome [18–21]. On the other hand, missense mutations in both the SAC1 and 5’-phosphophatase domains of *SYNJ1* have been found to associate with families of early onset atypical Parkinsonism with seizure [22–24]. The Parkinsonism linked R258Q mutation abolishes the SAC1 activity by ~80% [22, 25] and leads to dystrophic changes in both GABAergic and dopaminergic synapses in mice [25, 26]. The R839C mutation, which results in similar clinical phenotypes, has a more profound impact on both phosphatases as it reduces the 5’-phosphatase activity by ~60% and PI4P hydrolysis by 80% [25]. The functional relevance of the R839C mutation, however, hasn’t been explored.

Despite the well-known role of Synj1 in synaptic function, a few recent studies (including one from our lab) also suggest its involvement in autophagy regulation [12, 25]. Flies carrying the *Synj* R258Q mutation exhibit an impairment in autophagosome maturation at the neuromuscular junction (NMJ). In the aged *Synj1+/−* mouse brain lysate [25], we found an increase in lipidated LC3 (LC3-II), a hallmark of mature autophagosomes, as well as an increase in the autophagy substrate, p62, indicating an impairment in autolysosomal clearance. However, it was left unclear how astrocytes might have contributed to this pathology. Whether Synj1 regulates astrocyte function remains largely unknown, except for an earlier study that showed its potential contribution to astrogliogenesis [27]. Our current study using cultured astrocytes from *Synj1* littermate mice demonstrates that the neuronal isoform, Synj1-145 kDa, is also expressed in the astrocyte. More importantly, we show that endogenous Synj1 represses astrocyte autophagy at the basal level. *Synj1* deletion or the R839C mutation with a complex defect in the SAC1 and 5’-phosphatase activities leads to enhanced autophagosome formation and excessive autolysosomal clearance at the basal level. We demonstrate that Synj1’s role in phosphatidylinositol phosphate metabolism is important for maintaining a proper basal autophagy level, and that the enhanced basal autophagy in *Synj1* deficient conditions may be related to starvation at the resting state.

## Results

### Synj1 deletion induces astrocyte membrane stress

The role of Synj1 in astrocyte is poorly characterized. Using a Novus antibody that specifically recognizes the Synj1-145 kDa (Fig. 1a), we identified a low-level expression of Synj1 in the cultured cortical astrocytes, but not in microglia or HEK293T cells (Fig. S1a). Deletion of *Synj1* resulted in increased plasma membrane PI(4,5)P_2_, suggesting a prominent role of the 5’-phosphatase domain in astrocyte membrane signaling (Fig. S1b, c). To further investigate how Synj1 influences astrocyte membrane trafficking, we examined the endosomal markers, including EEA1 for early endosomes, Rab7 for late endosomes, and LAMP1 for lysosomes. Previous studies have shown that transgenic mice overexpressing Synj1 resulted in enlarged early endosome structures in neurons [20]; however, knocking down endogenous *Synj1* in Hela and SH-SY5Y cells also increased early endosomes without affecting the late endosomes [28].

**Fig. 1:**
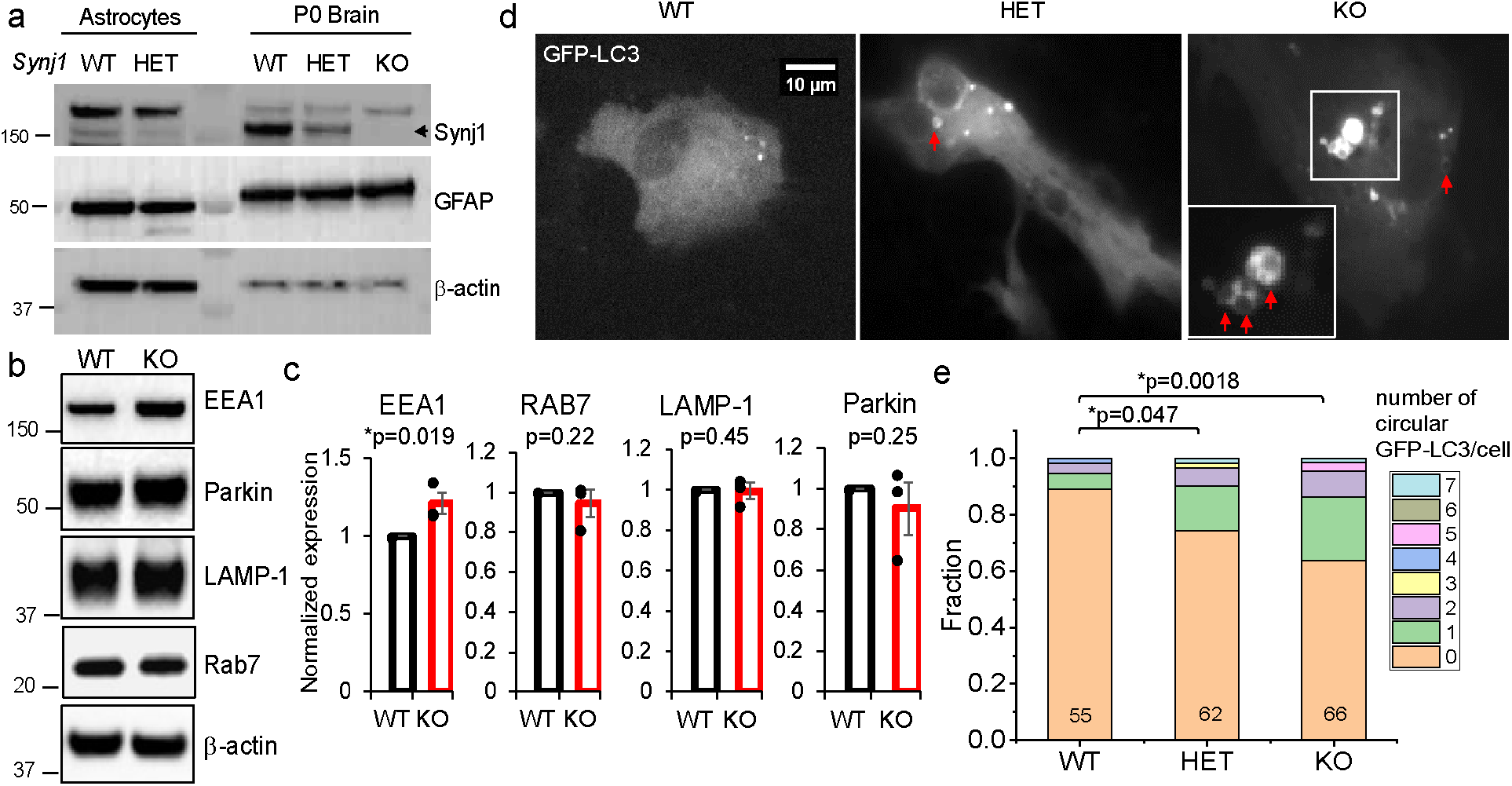
*Synj1* deletion induces astrocyte membrane stress. a) The NBP1-87842 rabbit anti-Synaptojanin1 from Novus Biologicals recognizes the 145 kDa isoform of Synj1 (a), which was abundantly expressed in the brain and weakly expressed in the astrocytes. b-c) Western blot analysis of the astrocyte lysate from *synj1* WT and KO littermates. Data from 3 independent batches of the astrocyte culture. P-values were from the Student’s *t*-test. d) Cultured astrocytes from *synj1* WT, HET and KO brains expressing GFP-LC3. Arrows point to circular LC3 structures. e) Analysis of the occurrence of circular GFP-LC3 structures in littermate astrocyte cultures presented by the stacked column plot. The number of cells analyzed from each genotype was indicated on the column. Data from 3 independent batches of culture. P-values were from the Mann Whitney *U* test.

In the present study examining *Synj1* knockout (KO) astrocytes, we observed a similar increase in the early endosome marker, EEA1 (Fig. 1b, c), while Rab7 and LAMP1 remained comparable to the wild-type (WT) cells. Parkin, the E3 ubiquitin ligase, which has an important role in mitophagy [29–31], was significantly upregulated in the *Synj1* R258Q knockin mice [26]. However, Parkin was not increased in *Synj1* KO astrocytes (Fig. 1b, c), suggesting that mitophagy is largely unaffected. Next, we expressed GFP-LC3 in cultured astrocytes to examine the autophagosomes, whose biogenesis is thought to rely on membrane sources including the plasma membrane and the endosomes [32]. In astrocytes prepared from *synj1+/+* (WT), *synj1+/−* (HET) and *synj1-/-* (KO) littermate pups, we observed LC3 puncta of varying sizes, some of which presented as circular structures with a hollow center that measured ~1-2 μm in diameter (Fig. 1d). Only a small fraction of these structures exhibited overlapping staining with Rab7 or EEA1 (Fig. S2), suggesting that they represent a distinct membrane compartment that is disorganized in *Synj1* deficient conditions. We quantified the number of these circular LC3 structures in all GFP-LC3 expressing astrocytes from 3 batches of primary littermate cultures and found a reverse gene dose-dependence in its occurrence. Our data suggests that Synj1 deficiency leads to membrane trafficking defects involving the endosomal compartments and the autophagosomes in astrocytes.

### Synj1 deficiency enhances the basal level autophagosome formation but impairs the starvation-induced autophagy clearance in astrocytes

Dysregulation of the autophagy pathway has been significantly implicated in multiple neurological disorders, including Parkinsonian neurodegeneration [1, 5, 33, 34]. A study by the Verstreken group showed that *Synj-/-* or the Parkinsonism linked *Synj* R258Q mutation impaired autophagosome maturation induced by starvation or prolonged stimulation at the fly NMJ [12]. In the *Synj1+/−* mouse, our previous study didn’t reveal any changes in LC3 immunofluorescence in the midbrain dopamine neurons, but found a reduction of p62 at 3 months of age with a robust increase at 18 months [25]. These results suggest an age-dependent reversal of the autolysosomal function in the dopamine neurons. Interestingly, although LC3 was not changed in the dopamine neurons of the *Synj1+/−* mice, an overall increase was noted in the striatum and the cortex, suggesting cell type-specific heterogeneity of the autophagy response to *Synj1* deficiency.

To elucidate the Synj1-mediated autophagy in astrocytes, we first analyzed the autophagy flux under basal conditions. The number of GFP-LC3 puncta in *Synj1* HET or KO astrocytes (HET: 6.95 ± 0.92, N= 57; KO: 8.46 ± 1.00, N= 91, 4 batches) was modestly, but significantly higher compared to that in the WT littermates (5.19 ± 0.73, N= 81, 4 batches) (Fig. 2a, c). In a separate set of littermate astrocytes, where bafilomycin A1 (baf, 20 nM) was applied for an hour to inhibit autolysosomal degradation, a much more striking increase was observed for *Synj1* deficient astrocytes (HET: 18 ± 2.54, N= 75 and KO: 26.50 ± 4.49, N= 58; 4 batches) compared to WT (8.74 ± 1.22, N= 62; 4 batches) (Fig. 2 a, d), consistent with our Western blot analysis (Fig. S3). Our results suggest hyperactive formation of autophagosome in *Synj1* deficient cells at the basal level. The autophagy substrate, p62, was hard to detect consistently in our Western blot analyses. However, consistent with the enhanced autophagosome formation, both *Synj1* HET and KO astrocytes exhibited a near 15% reduction in the p62 immunofluorescence compared to the WT cells expressing GFP-LC3 (Fig. 2b, e).

**Fig. 2:**
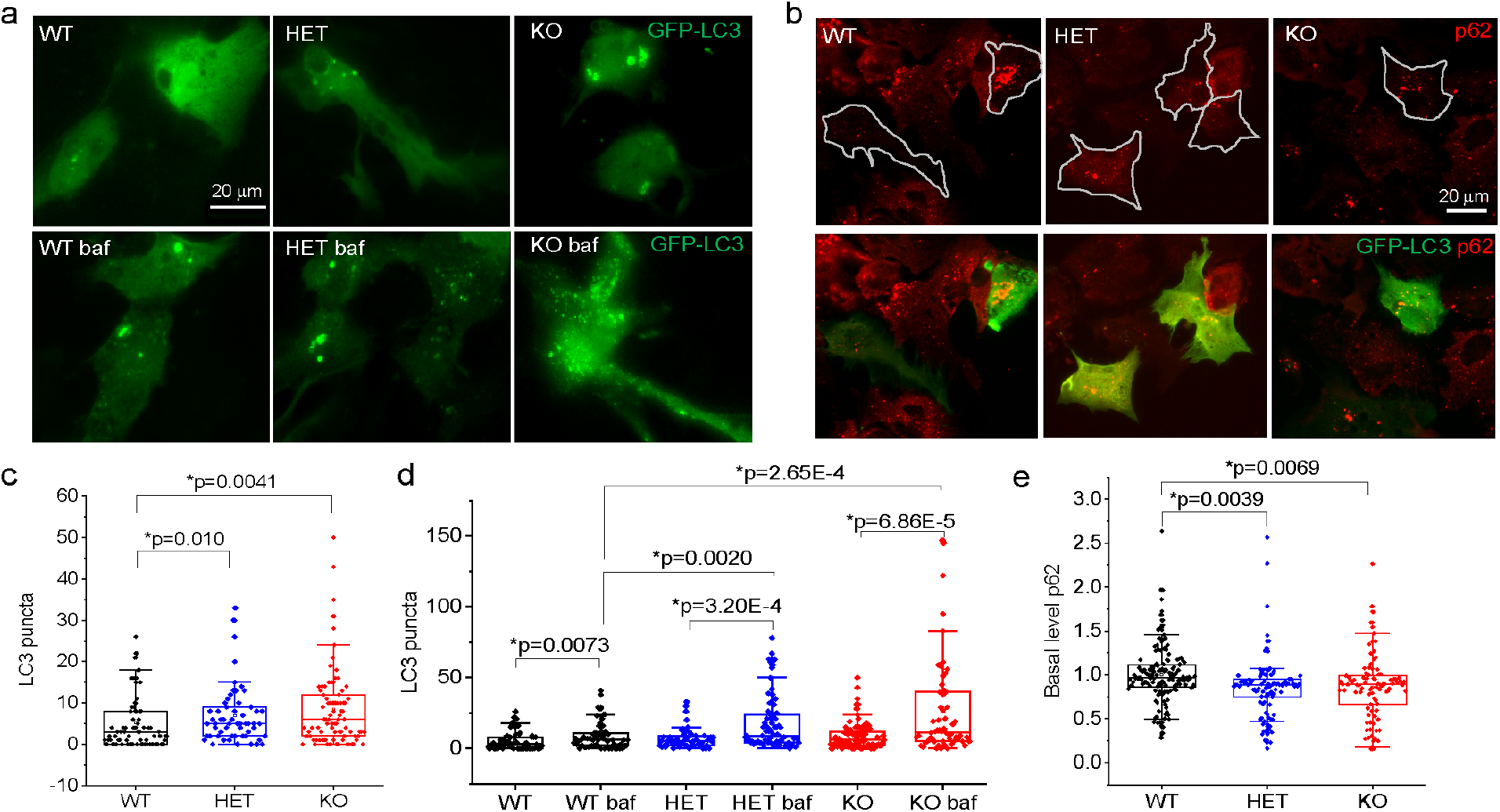
The basal level autophagosome formation is enhanced in *Synj1* deficient astrocytes. a-b) Representative images of cultured astrocytes from P0 *synj1* WT, HET and KO brains expressing GFP-LC3, those that treated with 20 nm bafilomycin A1 for 1 hour (lower panels in a) and those that immunolabeled with p62 (b). The white outlines in (b) indicate transfected cells that were included in the p62 analysis in (e). c-e) Box plots comparing the number of LC3 puncta at the basal level (c), those after the treatment of bafilomycin A1 (d), and p62 immunofluorescence in GFP-LC3 expressing cells (e). Data from 4 independent batches of cell cultures. P-values in (c) and (d) were from Mann-Whitney *U* tests and p-values in (e) were from Tukey’s *posthoc* following one-way ANOVA.

We next examined whether starvation-induced autophagosome formation or mTORC1 regulated autophagosome formation was affected by *Synj1* deficient conditions. In cultured WT neurons from the murine models, starvation or pharmacologically inhibiting the mTORC1 activity by rapamycin or torin has been shown as an ineffective means to induce autophagy [35]. Our previous study showed an intriguing sensitization of the *Synj1+/−* dopamine neurons to rapamycin-induced p62 clearance [25]. To examine the starvation induced autophagy in astrocytes, we used either a 4-hour rapamycin (200 nM) treatment, which inhibits the mTORC1 to induce autophagy [36, 37] or serum starvation (replacing the culture medium containing 10% FBS to the DMEM only medium) to mimic nutrient deprivation. For LC3 analysis, bafilomycin A1 was added in the last hour to reveal autophagy flux. Unlike the case in cultured neurons, we showed that WT astrocytes responded robustly and consistently to either rapamycin treatment or serum deprivation with an increase in GFP-LC3 puncta (control: 4.75 ± 0.53, N= 120; Rap: 22.72 ± 2.78, N= 82, DMEM: 23.22 ± 4.10, N= 70) (Fig. 3a, b) and a reduction in p62 immunofluorescence (normalized control: 1 ± 0.025, N= 109; normalized Rap: 0.83 ± 0.031, N= 82; normalized DMEM: 0.76 ± 0.041, N= 57) (Fig. 3c, d) in all 4 batches of cultures examined. The rapamycin responses were still present for LC3 in *Synj1* HET and KO astrocytes but significantly weaker (HET Rap: 18.49 ± 3.72, N= 69; KO Rap: 14.89 ± 2.08, N= 85) compared to the WT cells. Interestingly, unlike what we have found for *Synj1* deficient dopamine neurons, rapamycin was ineffective in clearing the autophagy substrate, p62, in both HET and KO astrocytes. Despite the weaker LC3 responses to rapamycin in *Synj1* deficient astrocytes, their responses to the 4-hour serum deprivation were as robust as the WT cells (KO DMEM: 28.41 ± 4.52, N= 78, HET DMEM: 21.10 ± 3.31, N= 50) (Fig. 3a, b, right panel). The p62 level, however, remained unaffected. Taken together, our data suggest that *Synj1* deficiency minimally impairs the starvation-induced autophagosome formation in astrocytes, but more dramatically decapacitates autolysosomal degradation under such conditions.

**Fig. 3:**
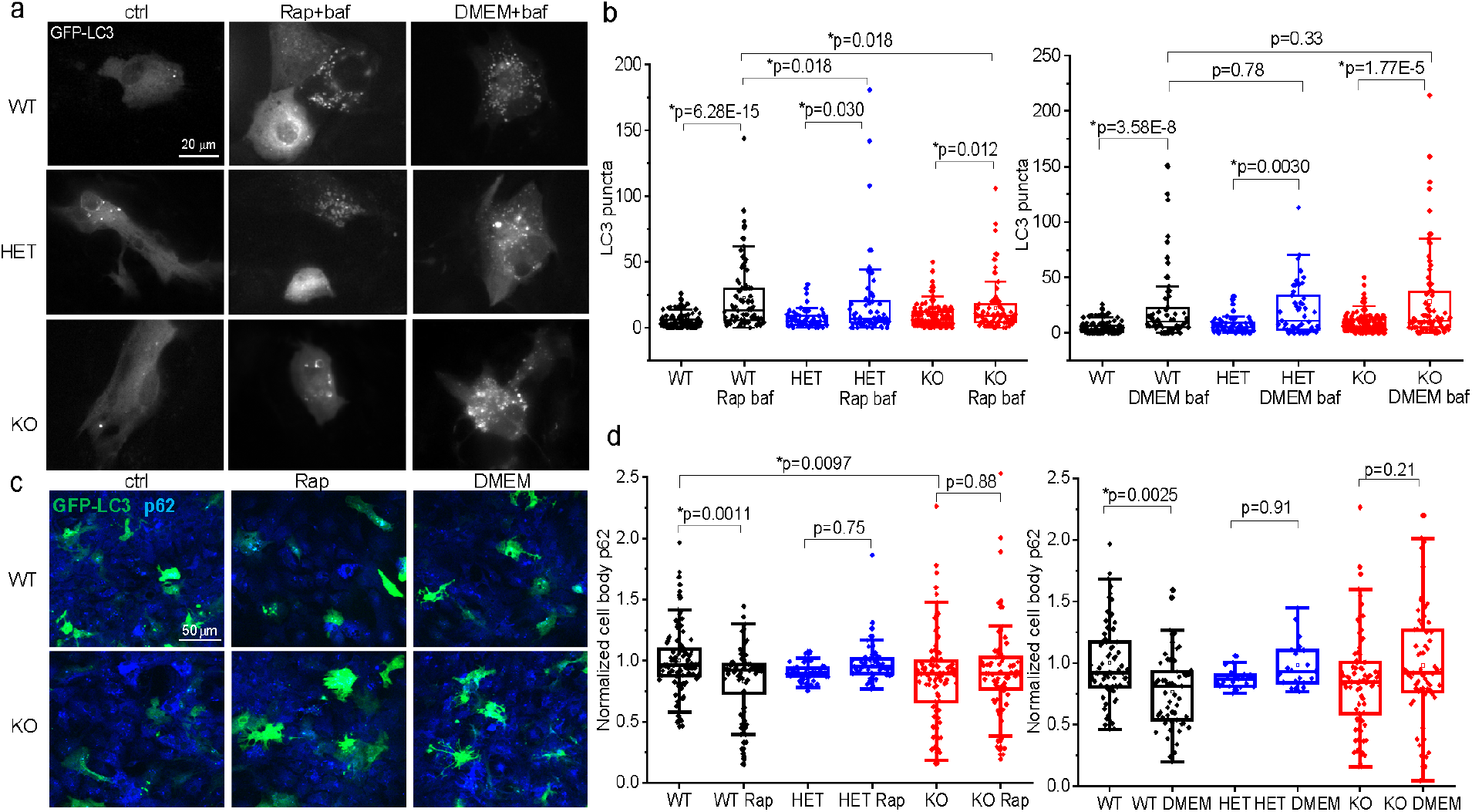
*Synj1* deficiency impairs starvation and rapamycin induced autophagy clearance. a) Representative images of cultured astrocytes from P0 *synj1* WT, HET and KO brains expressing GFP-LC3 (ctrl), those that were treated with 200 nM rapamycin (Rap+baf) or the DMEM-only medium (DMEM+baf) for 4 hours including a 20 nm bafilomycin A1 treatment in the last hour. b) Box plots comparing the number of LC3 puncta between the ctrl and the rapamycin treated groups (left), or between ctrl and the DMEM-only medium treated groups (right). P-values were from Mann-Whitney *U* tests. c) Representative images of cultured astrocytes from P0 *synj1* WT and KO brains expressing GFP-LC3 (ctrl), those that were treated with 200 nM rapamycin for 4 hours (Rap) and those that were treated with the DMEM-only medium for 4 hours. Cells were fixed simultaneously and immunolabeled with GFP and p62. d) Box plots comparing the normalized p62 levels across in the ctrl and the rapamycin treated groups (left), or between ctrl and the DMEM-only medium treated groups (right). P-values were from Tukey’s *posthoc* following two-way ANOVA. Data from 4 independent batches of cell cultures.

### The Synj1 phosphatase domains play a major role in repressing astrocyte basal level autophagy

Multiple Parkinsonism related *Synj1* mutations have been identified [22–24, 38] and all mutations reside in the two phosphatase domains (Fig. 4a). Our previous study using an *in vitro* phosphatase assay showed that the disease linked R258Q (RQ) mutation abolishes the PI4P and PI3P hydrolysis by ~80% [22, 25] while the R839C (RC) mutation reduces the 5’-phosphatase activity by ~60% and PI4P hydrolysis by 80%[25]. To date, studies of the RQ mutation suggest its role in maintaining axonal and synaptic morphology [26], autophagosome maturation, [12] and endosomal trafficking [28]; however, the functional relevance of the RC mutation hasn’t been documented.

**Fig. 4.**
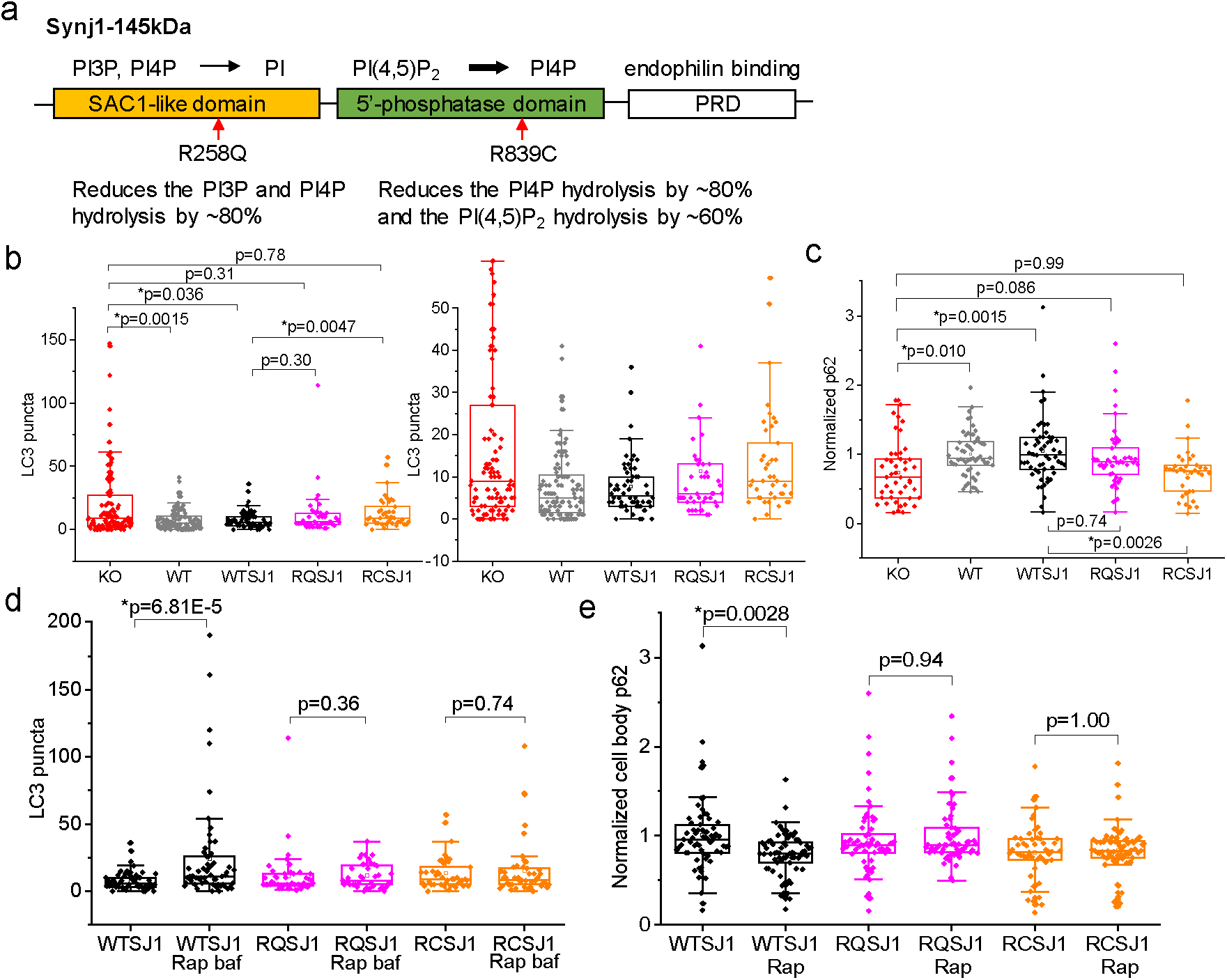
The Synj1 phosphatase domains play a major role in repressing astrocyte basal autophagy clearance. a) Domain structure and function of Synj1 with arrows pointing to the known Parkinsonism mutations illustrated by the functional outcome of the mutations reported in our previous publication [25]. b-e) WT hSynj1 (WTSJ1), R258Q hSynj1 (RQSJ1) or R839C hSynj1 (RCSJ1) were expressed in the *Synj1* KO astrocytes and compared to the KO and its littermate WT astrocyte culture. b-c) Basal level analysis: box plots comparing the number of LC3 puncta with 1-hour baf treatment for all groups (b) or the normalized p62 immunofluorescence (c) in the GFP-LC3 expressing astrocytes in different groups as indicated. d-e) Autophagy induction analysis: box plots comparing the number of GFP-LC3 puncta between ctrl and Rap+baf treatment groups as described in Fig. 3 (d), or the normalized p62 immunofluorescence in the ctrl and the rapamycin treated groups (e). P-values for the LC3 analyses were from Mann-Whitney *U* tests. P-values for p62 analyses were from Tukey’s *posthoc* tests following one-way ANOVA (c) or two-way ANOVA (e).

We expressed either the RQ hSynj1-145 kDa (RQ SJ1), RC hSynj1-145 kDa (RC SJ1) or WT hSynj1-145 kDa (WT SJ1) into the *Synj1* KO astrocyte culture and compared the basal level autophagy with the littermate WT culture and the KO culture. We found that the WT SJ1 expression effectively reversed the increased GFP-LC3 puncta number in KO astrocytes (KO: 20.78 ± 2.85, N= 105 compared to WT SJ1 rescue: 7.69 ± 0.90, N= 58 and WT littermate: 7.81± 0.79, N= 112; 3 batches) (Fig. 4b). However, the RC SJ1 was unable to repress the increased autophagosomes in the KO background (RC SJ1: 13.44 ± 1.94, N= 41, compared to KO: 20.78 ± 2.85, N= 105, p= 0.78 M-W *U* test) (Fig. 4b). Consistently, while the WT SJ1 effectively rescued the p62 level (KO normalized to WT littermate: 0.74 ± 0.064, N= 46 compared to WT SJ1 rescue normalized to WT littermate: 1.05 ± 0.062, N= 58 and WT littermate: 1.00± 0.042, N= 58; 3 batches), the RC SJ1 failed to do the same (RC SJ1 normalized to WT littermate: 0.73 ± 0.056, N= 36 compared to KO normalized to WT littermate: 0.74 ± 0.064, N= 46; 3 batches) (Fig. 4c). The rescue efficiency for the RQ SJ1 was in between the WT SJ1 and the RC SJ1. The LC3 puncta numbers and the p62 levels were neither different from the KO cells nor the WT SJ1 rescue (Fig. 4b, c). Our data suggest that both phosphatase activities of Synj1 are important for regulating basal level autophagy, but the RC mutation with a more profound defect in the phosphatase activities could produce a phenotype that mimics *Synj1* deletion.

We next examined how the Synj1 disease mutants affected the mTORC1 regulated autophagy. Not surprisingly, WT SJ1 was able to restore the rapamycin sensitivity in autophagosome formation (WT SJ1: 7.69 ± 0.90, N= 58; WT SJ1 Rap: 24.29 ± 4.78, N= 59; 3 batches) and p62 clearance (WT SJ1: 1.00 ± 0.053, N= 66; WT SJ1 Rap normalized to WT SJ1: 0.79 ± 0.029, N=70; 3 batches), but neither of the SJ1 mutants did (Fig. 4d, e). Due to concerns of SJ1 overexpression in these experiments, we performed a correlation analysis for Synj1 immunofluorescence (IF) and GFP-LC3 puncta as well as for Synj1 IF and p62 IF. Our data showed that in the subset of cells where the exogenous WT SJ1 was expressed at the endogenous level (determined by the WT littermate culture), the rescue for both GFP-LC3 and p62 was equally effective (Fig. S4). Overexpressing WT SJ1 did not lead to significant changes in either the LC3 puncta numbers or the p62 levels, nor did the SJ1 mutants.

### The energy sensing pathways are altered in Synj1-deficient astrocytes

To understand the molecular pathways that may underlie the enhanced basal level autophagy in Synj1 deficient astrocytes, we performed Western blot analysis. In nutrient rich conditions, autophagy is inhibited by mTOR (the mammalian/mechanistic target of rapamycin), a serine/threonine kinase essential in sensing energy metabolism and growth factors for coordinating cell growth [36]. Under starvation conditions, an imbalance of the AMP to ATP ratio activates the AMPK (AMP activated protein kinase) [39, 40], which can either directly induce autophagy or enhance autophagy through inhibiting the mTOR complex 1 (mTORC1) (Fig. 5a). We found that mTOR autophosphorylation at Ser2481 [41] was reduced by 50% in *Synj1* KO astrocytes compared to the WT littermates (Fig. 5b, c). The mTORC1 activity at Ser2448 was, however, not different between the WT and KO astrocytes at the basal level (Fig. 5d, e), despite an effective reduction upon rapamycin treatment and serum starvation (Fig. 5d). Interestingly, the ULK1/ATG1 expression was upregulated at the basal level in the KO astrocytes and the phosphorylation of ULK1 at Ser757, a direct substrate of the mTORC1 [42], was proportionally reduced (Fig. 5e), which may permit AMPK activated autophagy induction [42]. Indeed, AMPK phosphorylation at Thr172 was significantly enhanced in the KO astrocytes at the basal level (Fig. 5d, e), supporting the notion that *Synj1* deficient astrocytes suffer from nutrient deprivation at the basal level, which results in active autophagosome formation via deregulating both the AMPK and mTOR signaling pathways.

**Fig. 5.**
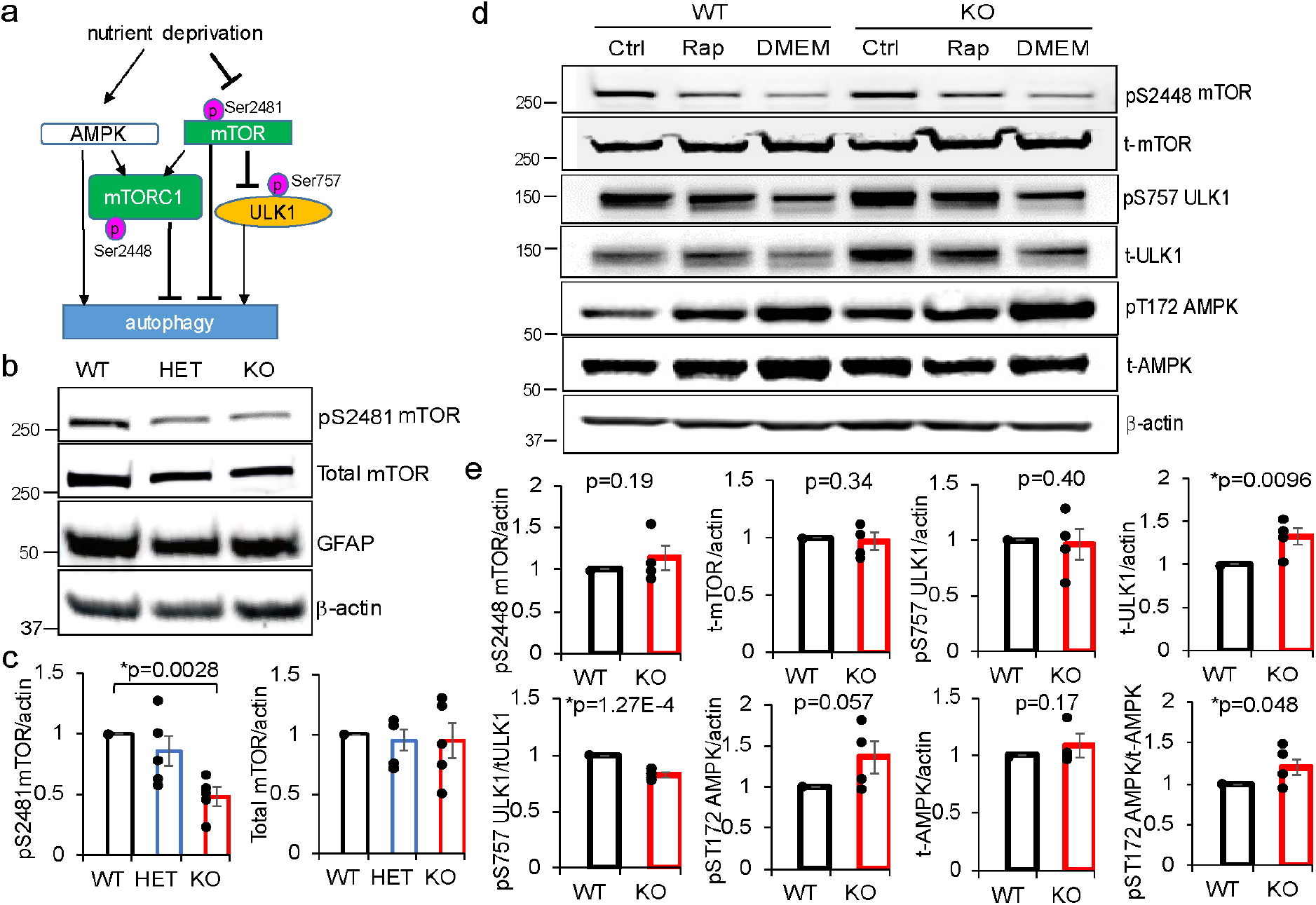
The energy sensing pathways are altered in *Synj1* deficient astrocytes. a) Signaling pathways that may influence the basal level astrocyte autophagy. b-c) Western blot analysis (b) and summary plots (c) for pS2481 mTOR and total mTOR expression in astrocyte cultures made from *Synj1* WT, HET and KO littermate pups. P-value was from Tukey’s *posthoc* following one-way ANOVA. d) Western blot analysis for mTORC1, ULK1 and AMPK activity in astrocyte cultures made from *Synj1* WT and KO littermate pups treated with rapamycin or DMEM as described in Fig. 3. e) Summary of the pS2448 mTOR, pS757 ULK1 and pT172 AMPK activity and their protein expression at the basal level from 4 independent batch of cultures. P-values were from Student’s *t*-test.

## Discussion

Our study demonstrates that the well-characterized neuronal protein, Synj1, is also present in astrocytes. We further reveal a novel role for Synj1 in repressing astrocyte autophagy at the basal level. We show that the phosphatase activities of Synj1 are important in keeping the basal level autophagy in check. The Parkinsonism linked R839C mutation, which has a more profound impact on the phosphatidylinositol phosphate metabolism than the R258Q mutation [25], is reported for the first time to promote basal level autophagosome formation and clearance that mimicked *Synj1* deletion. We found that although the basal level autophagy was enhanced in *Synj1* deficient astrocytes, the starvation/rapamycin induced autolysosomal degradation was impaired. Neither the R258Q nor the R839C mutation rescued the rapamycin induced autophagy defect in *Synj1* KO astrocytes, suggesting that the intact phosphatase activities in Synj1 are especially crucial for astrocytes to cope with cellular stress such as nutrient starvation.

Additional molecular machineries may be involved during stress induced autophagy. In an earlier study, it was suggested that PI(3,5)P_2_ accumulation due to the R258Q mutation could impede LC3 lipidation in the drosophila NMJ [12]. However, an enhanced basal level autophagy was not reported in the *Synj* mutant drosophila, although it was reminiscent of our previous findings in the *Synj1+/−* mouse brains [25], where we showed an increase in the endogenous LC3 immunofluorescence in the cortex and striatum. While we were unable to distinguish an effect from the glia or the neuron then, our current study suggests that the increased LC3 might in part be due to an upregulated autophagy pathway in the astrocytes at the basal level.

Most strikingly, we show that at the basal level, a 1-hour bafilomycin A1 treatment for the KO astrocytes resulted in a 3-fold increase in the GFP-LC3 puncta (average 26.50 ± 4.49 puncta/cell, Fig. 2c), an effect as potent as the WT astrocytes with a 4-hour rapamycin treatment or serum deprivation (Rap: 22.72 ± 2.78, DMEM: 23.22 ± 4.10, Fig. 3b). Our data suggests that at the basal level, a remarkable portion of the energy supply for the KO astrocytes might be coming from lysosomal degradation, a process suggested by earlier studies as an adaptive response to prolonged starvation [43–46]. When bafilomycin was applied to inhibit lysosomal degradation, autophagy was induced via inhibiting the lysosome targeted mTORC1. The energy supply is likely different in the WT astrocytes, whose nutrients come mostly from external sources, such as the serum. Therefore, rapamycin or serum deprivation resulted in a much more robust autophagy response compared to the bafilomycin treatment. These data were consistent with our biochemical finding that the KO astrocytes that were starved at the basal level exhibited reduced mTOR and increased AMPK activities (Fig. 5).

The inability to source nutrients from the external environment could lead to an adaptive upregulation of basal level autophagosome formation. It remains to be investigated why *Synj1* deficiency impaired the cell’s capability to source external nutrients. A previous report showed that the glucose transporter (GluT1) level was reduced in the *Synj1-/-* mice [27]. The result, however, was not replicated in our astrocyte cultures derived from the *Synj1-/-* brains (data not shown). Nonetheless, possibilities remain that *Synj1* deficiency affects the surface expression of GluT1, or amino acid transporters on the plasma membrane [47–49]. Our data suggests that the phosphatase activities of Synj1 are likely responsible for setting the energy state of the astrocytes. The Parkinsonism R839C mutation that disrupts both phosphatase domains can enhance autophagic clearance at the basal level, mimicking that of *Synj1* KO astrocytes. While the R258Q mutation that abolishes the SAC1 activity did little to the basal level autophagy, it remains to be clarified if the accumulation of PI(4,5)P_2_ is the culprit for astrocyte energy sensing.

Interestingly, while *Synj1* deficiency didn’t affect lysosomal digestion at the basal level (Fig. 2e), the rapamycin or serum deprivation induced autolysosomal degradation was significantly impaired (Fig. 3d). The defect could be due to an intrinsic dysfunction of the lysosomal system under stress, or the already elevated lysosomal activity at the baseline, which leaves little capacity for the astrocytes to handle additional workload. The exuberant self-digestion at the baseline, along with the inflexibility of the astrocytes to respond to environmental stressors, such as starvation, could be detrimental to cell survival [50].

As multiple missense mutations in *SYNJ1* were found in Parkinsonian patients with seizure, it remains to be investigated whether and how dysfunctional astrocyte autophagy contributes to the pathogenic course *in vivo*. Most importantly, will the exuberant phagocytic activity in astrocytes result in dopaminergic synapse pruning [51]? Or will the inflexibility of astrocytes to respond to external stressors result in an accumulation of alpha-synuclein [25] and reactive oxygen species? Astrocytes, as major supporting cells in the brain, are actively involved in secretion of neurotrophic factors and cytokines to regulate neuronal plasticity and neuroinflammation [52, 53]. Astrocytes from different parts of the mouse brain have also been shown to perform different roles to support the neuron [54, 55]. It remains to be researched if astrocytes from the striatum or the midbrain also exhibit altered autophagy function to ultimately impact dopaminergic neurotransmission and neuronal survival [56].

Despite the conserved pool of autophagy genes from yeast to mammalian cells, tremendous differences exist across species and cell types for its regulatory mechanisms [5, 35, 57, 58]. Although neuronal autophagy has been shown to be less influenced by nutrient deficiency [35, 59, 60], it remains to be elucidated if an enhanced basal level autophagy flux is also present in *Synj1* deficient murine and human neurons. Thus, our study reveals a novel role of Synj1 in astrocyte autophagy/energy sensing, and brings insight to Synj1-mediated pathogenic mechanisms.

## Experimental Procedures

### Animals

Mice were housed in the pathogen-free barrier facility at the Rutgers Robert Wood Johnson Medical School SPH vivarium. Handling procedures were in accordance with the National Institutes of Health guidelines approved by the institutional Animal Care and Use Committee (IACUC). The *Synj1+/−* mice [9] were obtained from the Pietro De Camilli laboratory at Yale University. As the *Synj1-/-* mouse is lethal at birth, *Synj1+/−* mice were used as breeders to generate knockout (KO) pups and littermates.

### Cell culture and transfection

Astrocyte cultures were prepared from postnatal day 0 (P0) littermate pups of both sexes using a slightly modified protocol from published methods [61]. Mice were decapitated by sharp scissors and the brains were dissected in ice cold Hank’s solution (Sigma H2387) containing 350 mg/L NaHCO_3_ and 1 mM HEPES (260mg/L) with pH adjusted to 7.15~7.20. Typically, 2 cortices from mice of each genotype were dissected and broken into smaller pieces by the spring scissors. The tissues were then digested in room temperature for 7 min in a 3-mL Hank’s solution containing 0.25% Trypsin (Thermo Fisher, 15090046) and 0.1 μg/μL DNase (Sigma D5025) with intermittent shaking. Tissues were mechanically dissociated by pipetting and the trypsin reaction was terminated by addition of 4 mL culture media containing DMEM (Thermo Fisher, 11965118), 10% Fetal Bovine Serum (Atlantic Biologicals, S11550) and 10 U/mL Penicillin-Streptomycin (Thermo Fisher, 15140122). Cells were centrifuged at 300 g for 10 min and plated at ~8,000,000/10 cm dish pre-coated with poly-D-lysine (Sigma # A-003-E, 0.1 mg/mL). Culture medium was changed every 2-3 days after plating and cells typically reach 90% confluency after 10 days. To obtain an enriched astrocyte culture, the culture dish was placed on an orbital shaker at ~180 rpm for 30 min to remove microglia (or to obtain a separate microglia culture) and an additional 6 hours at ~240 rpm to remove oligodendrocytes precursor cells [61]. The remaining confluent astrocyte culture was rinsed by PBS and digested with 0.05% trypsin-EDTA (Thermo Fisher, 25300054). Enriched astrocytes from the first or the second passage were grown on cover glasses (#1.5) for imaging studies or on 6-well plates for Western blot analysis. HEK293T cells were grown in the same culture media as the astrocyte culture and maintained/passaged using the same procedure. For imaging analysis, cells were plated at 50% confluency and transfected the next day with GFP-LC3 (Addgene #21073) or double transfected with GFP-LC3 and one of the FLAG-hSynj1 constructs (see below). The Lipofectamine3000 reagent was used for transfection following a company suggested protocol.

### Constructs

pEGFPC1-FLAG-WT *hSYNJ1-*145 kDa, pEGFPC1-FLAG-R258Q *hSYNJ1-*145 kDa and pEGFPC1-FLAG-R839C *hSYNJ1-*145 kDa [25] were re-engineered by site directed mutagenesis (Agilent QuikChange 200517) to delete the EGFP using the following primers: 5’-CGCTAGCGCTACCGGTCGCCACCTCCGGACTCAGATC-3’ and 5’-GCTTGAGCTCGAGATCTGAGTCCGGAGGTGGCGACCGG-3’. The successful deletion of the EGFP was verified by sequencing.

### Western blot analysis and antibodies

Brain samples or cells were lysed on ice for 30 min using a Triton-based lysis buffer containing 50 mM Tris-HCl (pH 7.5), 150 mM NaCl, 1% Triton as well as protease and phosphatase inhibitors as previously described [16, 25]. After centrifugation at 16,000 g, 4°C for 30 min, supernatant was collected for protein quantification using the Pierce BCA assay (Thermo 23227). Typically, 5-10 μg of total proteins were loaded for each sample on the Invitrogen 4-12% Bis-Tris gel and the following antibodies were used for immunoblot detection: rabbit anti-Synj1 (Novus Biologicals, NBP1-87842, 1:1000), rabbit anti-GFAP (Abclonal, A0237, 1:2000), rabbit anti-EEA1 (Abcam, ab109110, 1:5000), mouse anti-Rab7 (Abcam, ab50533, 1:1000), rabbit anti-LAMP1 (Abcam, ab24170, 1:1000), mouse anti-Parkin (Cell signaling, 42115, 1:1000), rabbit anti-pS2481 mTOR (Cell signaling, 2974, 1:1000), rabbit anti-pS2448 mTOR (Cell signaling, 2971, 1:1000), rabbit anti-mTOR (Cell signaling, 2983, 1:1000), rabbit anti-pS757 ULK1 (Cell signaling, 14202, 1:1000), rabbit anti-ULK1 (Cell signaling, 6439, 1:1000), rabbit anti-pT172 AMPK (Cell signaling, 2535, 1:1000), rabbit anti-AMPKα (Cell signaling, 2532, 1:1000), rabbit anti-GLUT1 (Abcam, ab115730, 1:1000), mouse anti-β-actin (Cell signaling, 37005, 1:3000). All Western blots were performed with 2-3 technical repeats and the Western blot bands were analyzed using imageJ.

### Immunofluorescence

analysis -The following antibodies were used for immunofluorescence: mouse anti-PI(4,5)P_2_ (Echelon Biosciences, z-P045, 1:100) [25], chicken anti-GFP (Thermo Fisher, A-10262, 1:1000), guinea pig anti-p62 (Progene, GPP62-C, 1:1000), rabbit anti-EEA1 (Abcam, ab109110, 1:1000), mouse anti-Rab7 (Abcam, ab50533, 1:200). Immunocytochemistry was performed following previously published procedures [16, 25, 62]. Immuofluorescence was analyzed using a Nikon Ti-2 wide-field microscope with Spectra-X (Lumencor) as the light source and an Andor Ultra 897 EMCCD camera. The Alexa fluo-488, Alexa fluo-546 and Alexa fluo-647 emissions were collected using the ET535/50m, ET585/40m and ET665lp emission filters, respectively. All imaging parameters were set to the same for each batch of culture. Image stacks were taken at different focal planes at 0.9 μm interval to include the whole cell and a maximum projection image was generated for each stack via ImageJ for analysis. All analyses were done manually. The GFP-LC3 punctum was determined by 1.5 × 1.5 μm (6 × 6 pixels) circular regions of interests (ROIs), which means larger punctum was counted as multiple puncta. The p62 level was analyzed as whole cell immunofluorescence in randomly transfected GFP-LC3 cells.

### Data analysis

All imaging and Western blot studies were from at least 3 independent primary cultures. Most Western blots were repeated 3 times. The number of GFP-LC3 puncta at the basal level remained consistent in all batches. The p62 immunofluorescence and protein expression levels in Western blots varied across different batches and were all normalized to the values obtained in the WT cells in the same batch. The LC3 quantification data does not follow normal distribution and the statistical difference was calculated using Mann-Whitney *U* test. For datasets following normal distribution, Student’s *t*-test or One-way/Two-way ANOVA was performed followed by Tukey’s *posthoc* analysis using the build-in functions in OriginLab.

## Data availability

All data and plasmids described in the manuscript are available for sharing upon request.

## Acknowledgement

We thank Dr. Wei-xing Zong for discussion of the work and suggestions for the manuscript preparation.

## Funding and additional information

The work was funded by the NINDS R01 grant (R01NS112390) to P-Y Pan. The content is solely the responsibility of the authors and does not necessarily represent the official views of the National Institutes of Health.

---

**Fig. S1:**
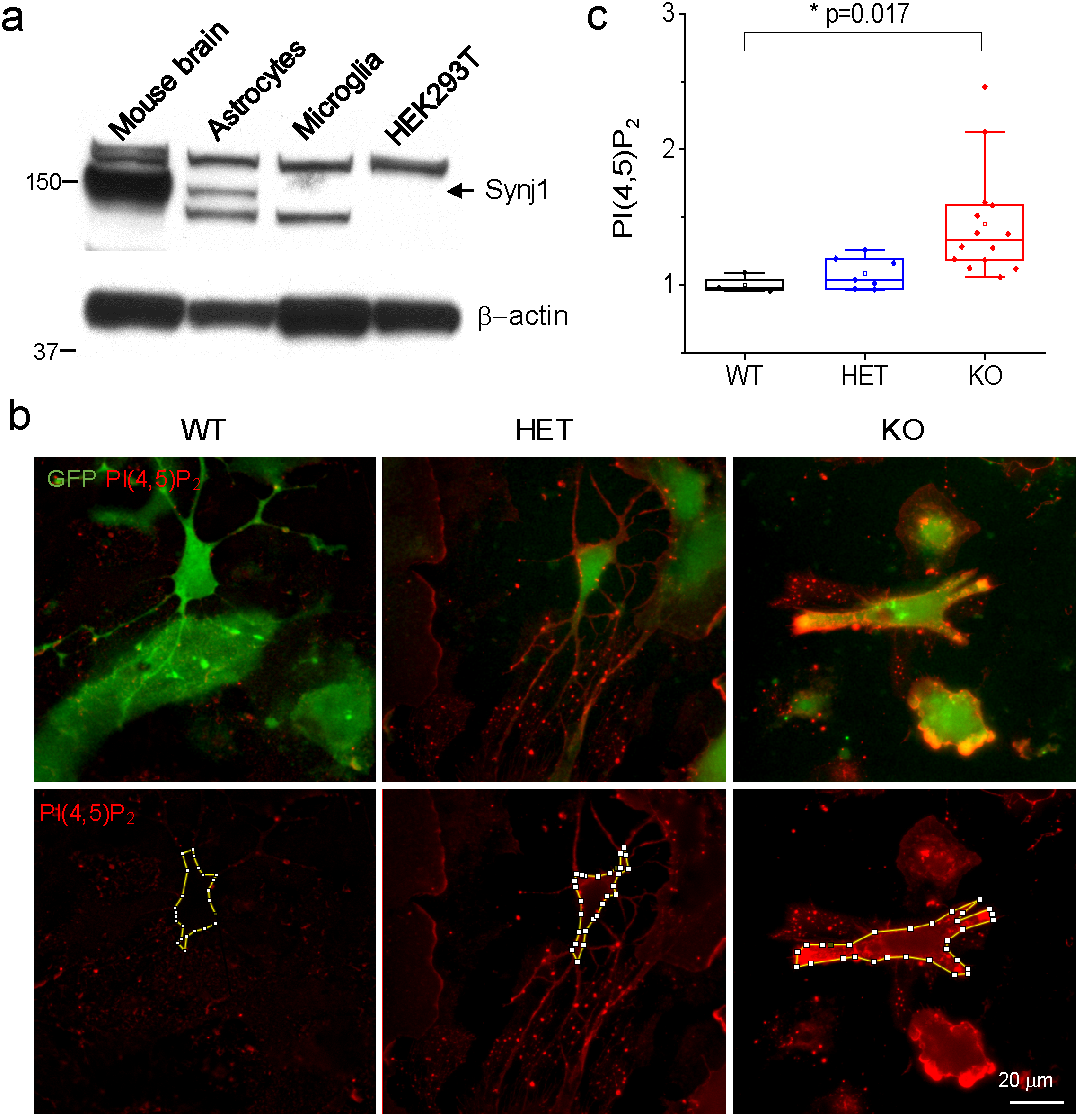
*Synj1* deletion in astrocytes results in increased membrane PI(4,5)P_2_. a) Western blot analysis of Synj1 expression in adult mouse brain lysate, astrocyte lysate, microglia lysate and HEK297T cell lysate as indicated using the NBP1-87842 polyclonal antibody. b) Immunofluorescence for PI(4, 5)P_2_ in *Synj1* WT, HET and KO astrocytes sparsely expressing GFP-LC3. The PI(4, 5)P_2_ antibody was validated in our previous report [25]. The yellow segmented lines were membrane selections used for PI(4, 5)P_2_ analysis. c) Analysis of the membrane PI(4, 5)P_2_ by tracing the contour of the transfected astrocytes shown in (b). P-value was from Tukey’s *posthoc* following one-way ANOVA.

**Fig. S2.**
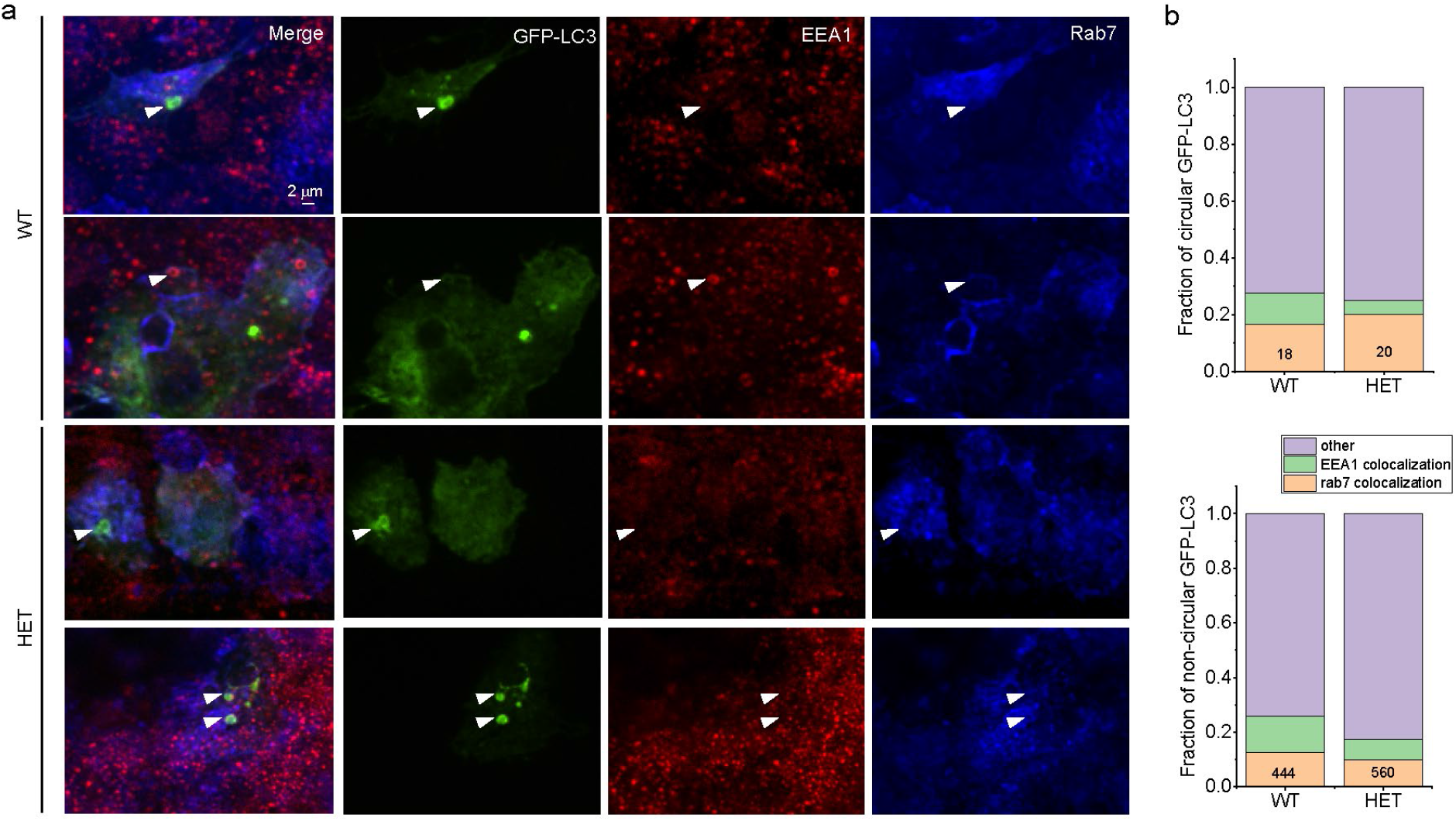
Few circular GFP-LC3 structures are colocalized with EEA1 or Rab7. a) WT and HET astrocytes were transfected with GFP-LC3 and fixed for immunolabeling. Arrow heads point to circular GFP-LC3 structures in WT and HET astrocytes. b) A small fraction of circular GFP-LC3 and non-circular GFP-LC3 structures were colocalized with EEA1 or Rab7. The total numbers of GFP-LC3 structures in each category were indicated.

**Fig. S3:**
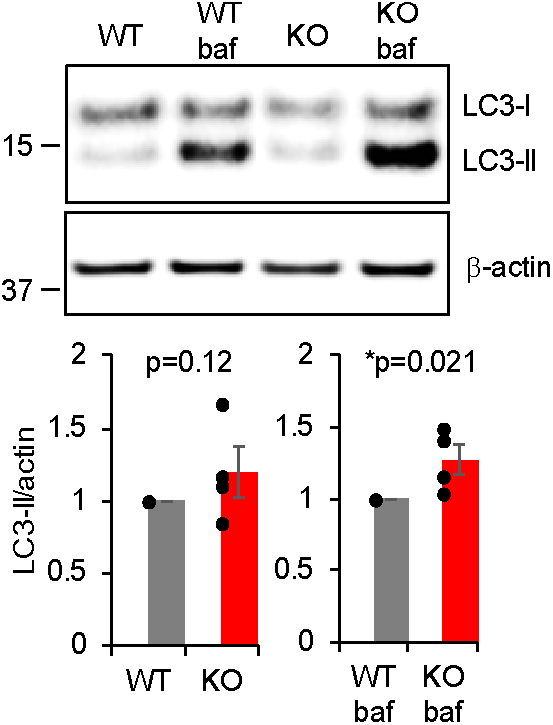
Bafilomycin treatment reveals increased LC3 lipidation in *Synj1* KO astrocytes. Western blot analysis for cultured WT and KO astrocytes at baseline and those that were treated with bafilomycin A1 for 1 hour. Data from 4 batches of cells and p-values were from Student’s *t*-test.

**Fig. S4:**
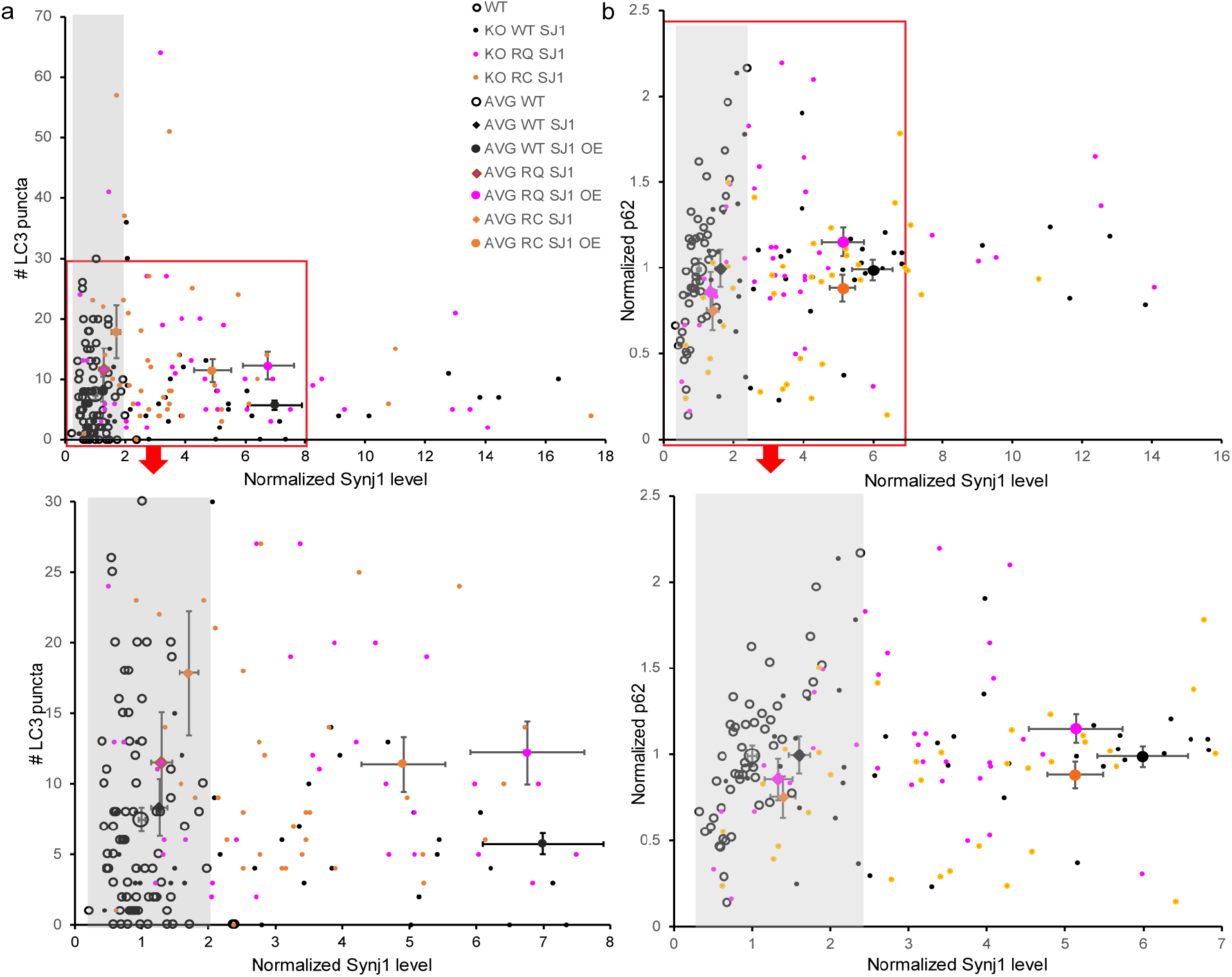
Synj1 overexpression does not affect astrocyte autophagy. a, b) Two-dimensional plot for the correlation between the number of GFP-LC3 puncta and the Synj1 expression level (a) or the p62 level and the Synj1 expression level (b). Data were extrapolated from Fig. 4. Synj1 expression levels measured in the WT cells were considered physiological levels of Synj1 (normalized to 1) with ranges indicated by the shaded grey boxes. The highest physiological level of Synj1 in WT cells divides each dataset (either expressing WT, RQ or RC synj1) into a rescue/replacement group (KO WT SJ1, KO RQ SJ1 or KO RC SJ1) and an overexpression (OE) group WT SJ1 OE, RQ SJ1 OE, or RC SJ1 OE). Each small symbol represents a measurement from a cell, and large symbols with error bars were averages from either the rescue/replacement or the OE group. Two-way ANOVA for protein expression and genotype followed by *posthoc* didn’t reveal any significant difference. Neither did the correlation tests.

## References

1. Nixon, R.A., The role of autophagy in neurodegenerative disease. Nat Med, 2013. 19(8): p. 983–97.

2. Menzies, F.M., A. Fleming, and D.C. Rubinsztein, Compromised autophagy and neurodegenerative diseases. Nat Rev Neurosci, 2015. 16(6): p. 345–57.

3. Lee, H.J., et al., Direct transfer of alpha-synuclein from neuron to astroglia causes inflammatory responses in synucleinopathies. J Biol Chem, 2010. 285(12): p. 9262–72.

4. Lindstrom, V., et al., Extensive uptake of alpha-synuclein oligomers in astrocytes results in sustained intracellular deposits and mitochondrial damage. Mol Cell Neurosci, 2017. 82: p. 143–156.

5. Tremblay, M.E., M.R. Cookson, and L. Civiero, Glial phagocytic clearance in Parkinson’s disease. Mol Neurodegener, 2019. 14(1): p. 16.

6. Sung, K. and M. Jimenez-Sanchez, Autophagy in Astrocytes and its Implications in Neurodegeneration. J Mol Biol, 2020. 432(8): p. 2605–2621.

7. McPherson, P.S., et al., A presynaptic inositol-5-phosphatase. Nature, 1996. 379(6563): p. 353–7.

8. Haucke, V. and G. Di Paolo, Lipids and lipid modifications in the regulation of membrane traffic. Curr Opin Cell Biol, 2007. 19(4): p. 426–35.

9. Cremona, O., et al., Essential role of phosphoinositide metabolism in synaptic vesicle recycling. Cell, 1999. 99(2): p. 179–88.

10. Mani, M., et al., The dual phosphatase activity of synaptojanin1 is required for both efficient synaptic vesicle endocytosis and reavailability at nerve terminals. Neuron, 2007. 56(6): p. 1004–18.

11. Cao, M., et al., Absence of Sac2/INPP5F enhances the phenotype of a Parkinson’s disease mutation of synaptojanin 1. Proc Natl Acad Sci U S A, 2020. 117(22): p. 12428–12434.

12. Vanhauwaert, R., et al., The SAC1 domain in synaptojanin is required for autophagosome maturation at presynaptic terminals. EMBO J, 2017. 36(10): p. 1392–1411.

13. Micheva, K.D., B.K. Kay, and P.S. McPherson, Synaptojanin forms two separate complexes in the nerve terminal. Interactions with endophilin and amphiphysin. J Biol Chem, 1997. 272(43): p. 27239–45.

14. Lee, S.Y., et al., Regulation of synaptojanin 1 by cyclin-dependent kinase 5 at synapses. Proc Natl Acad Sci U S A, 2004. 101(2): p. 546–51.

15. Dong, Y., et al., Synaptojanin cooperates in vivo with endophilin through an unexpected mechanism. Elife, 2015. 4.

16. Pan, P.Y., et al., Parkinson’s Disease-Associated LRRK2 Hyperactive Kinase Mutant Disrupts Synaptic Vesicle Trafficking in Ventral Midbrain Neurons. J Neurosci, 2017. 37(47): p. 11366–11376.

17. Haffner, C., et al., Direct interaction of the 170 kDa isoform of synaptojanin 1 with clathrin and with the clathrin adaptor AP-2. Curr Biol, 2000. 10(8): p. 471–4.

18. Voronov, S.V., et al., Synaptojanin 1-linked phosphoinositide dyshomeostasis and cognitive deficits in mouse models of Down’s syndrome. Proc Natl Acad Sci U S A, 2008. 105(27): p. 9415–20.

19. Zhu, L., et al., Reduction of synaptojanin 1 accelerates Abeta clearance and attenuates cognitive deterioration in an Alzheimer mouse model. J Biol Chem, 2013. 288(44): p. 32050–63.

20. Cossec, J.C., et al., Trisomy for synaptojanin1 in Down syndrome is functionally linked to the enlargement of early endosomes. Hum Mol Genet, 2012. 21(14): p. 3156–72.

21. McIntire, L.B., et al., Reduction of synaptojanin 1 ameliorates synaptic and behavioral impairments in a mouse model of Alzheimer’s disease. J Neurosci, 2012. 32(44): p. 15271–6.

22. Krebs, C.E., et al., The Sac1 domain of SYNJ1 identified mutated in a family with early-onset progressive Parkinsonism with generalized seizures. Hum Mutat, 2013. 34(9): p. 1200–7.

23. Quadri, M., et al., Mutation in the SYNJ1 gene associated with autosomal recessive, early-onset Parkinsonism. Hum Mutat, 2013. 34(9): p. 1208–15.

24. Taghavi, S., et al., A Clinical and Molecular Genetic Study of 50 Families with Autosomal Recessive Parkinsonism Revealed Known and Novel Gene Mutations. Mol Neurobiol, 2018. 55(4): p. 3477–3489.

25. Pan, P.Y., et al., Synj1 haploinsufficiency causes dopamine neuron vulnerability and alpha-synuclein accumulation in mice. Hum Mol Genet, 2020. 29(14): p. 2300–2312.

26. Cao, M., et al., Parkinson Sac Domain Mutation in Synaptojanin 1 Impairs Clathrin Uncoating at Synapses and Triggers Dystrophic Changes in Dopaminergic Axons. Neuron, 2017. 93(4): p. 882–896 e5.

27. Herrera, F., et al., Synaptojanin-1 plays a key role in astrogliogenesis: possible relevance for Down’s syndrome. Cell Death Differ, 2009. 16(6): p. 910–20.

28. Fasano, D., et al., Alteration of endosomal trafficking is associated with early-onset parkinsonism caused by SYNJ1 mutations. Cell Death Dis, 2018. 9(3): p. 385.

29. Wong, Y.C. and E.L. Holzbaur, Optineurin is an autophagy receptor for damaged mitochondria in parkin-mediated mitophagy that is disrupted by an ALS-linked mutation. Proc Natl Acad Sci U S A, 2014. 111(42): p. E4439–48.

30. Pickrell, A.M., et al., Endogenous Parkin Preserves Dopaminergic Substantia Nigral Neurons following Mitochondrial DNA Mutagenic Stress. Neuron, 2015. 87(2): p. 371–81.

31. Sliter, D.A., et al., Parkin and PINK1 mitigate STING-induced inflammation. Nature, 2018. 561(7722): p. 258–262.

32. Tooze, S.A., Current views on the source of the autophagosome membrane. Essays Biochem, 2013. 55: p. 29–38.

33. Pan, P.Y., et al., Crosstalk between presynaptic trafficking and autophagy in Parkinson’s disease. Neurobiol Dis, 2019. 122: p. 64–71.

34. Karabiyik, C., M.J. Lee, and D.C. Rubinsztein, Autophagy impairment in Parkinson’s disease. Essays Biochem, 2017. 61(6): p. 711–720.

35. Maday, S. and E.L. Holzbaur, Compartment-Specific Regulation of Autophagy in Primary Neurons. J Neurosci, 2016. 36(22): p. 5933–45.

36. Saxton, R.A. and D.M. Sabatini, mTOR Signaling in Growth, Metabolism, and Disease. Cell, 2017. 168(6): p. 960–976.

37. Smith, E.D., et al., Rapamycin and interleukin-1beta impair brain-derived neurotrophic factor-dependent neuron survival by modulating autophagy. J Biol Chem, 2014. 289(30): p. 20615–29.

38. Kirola, L., et al., Identification of a novel homozygous mutation Arg459Pro in SYNJ1 gene of an Indian family with autosomal recessive juvenile Parkinsonism. Parkinsonism Relat Disord, 2016. 31: p. 124–128.

39. Cork, G.K., J. Thompson, and C. Slawson, Real Talk: The Inter-play Between the mTOR, AMPK, and Hexosamine Biosynthetic Pathways in Cell Signaling. Front Endocrinol (Lausanne), 2018. 9: p. 522.

40. Carling, D., The AMP-activated protein kinase cascade--a unifying system for energy control. Trends Biochem Sci, 2004. 29(1): p. 18–24.

41. Soliman, G.A., et al., mTOR Ser-2481 autophosphorylation monitors mTORC-specific catalytic activity and clarifies rapamycin mechanism of action. J Biol Chem, 2010. 285(11): p. 7866–79.

42. Kim, J., et al., AMPK and mTOR regulate autophagy through direct phosphorylation of Ulk1. Nat Cell Biol, 2011. 13(2): p. 132–41.

43. Yu, L., et al., Termination of autophagy and reformation of lysosomes regulated by mTOR. Nature, 2010. 465(7300): p. 942–6.

44. Zoncu, R., et al., mTORC1 senses lysosomal amino acids through an inside-out mechanism that requires the vacuolar H(+)-ATPase. Science, 2011. 334(6056): p. 678–83.

45. Yao, Y., E. Jones, and K. Inoki, Lysosomal Regulation of mTORC1 by Amino Acids in Mammalian Cells. Biomolecules, 2017. 7(3).

46. Guo, J.Y., et al., Autophagy provides metabolic substrates to maintain energy charge and nucleotide pools in Ras-driven lung cancer cells. Genes Dev, 2016. 30(15): p. 1704–17.

47. Pinilla, J., et al., SNAT2 transceptor signalling via mTOR: a role in cell growth and proliferation? Front Biosci (Elite Ed), 2011. 3: p. 1289–99.

48. Wang, Q., et al., Targeting amino acid transport in metastatic castration-resistant prostate cancer: effects on cell cycle, cell growth, and tumor development. J Natl Cancer Inst, 2013. 105(19): p. 1463–73.

49. Nicklin, P., et al., Bidirectional transport of amino acids regulates mTOR and autophagy. Cell, 2009. 136(3): p. 521–34.

50. Bialik, S., S.K. Dasari, and A. Kimchi, Autophagy-dependent cell death - where, how and why a cell eats itself to death. J Cell Sci, 2018. 131(18).

51. Lieberman, O.J., et al., Roles for neuronal and glial autophagy in synaptic pruning during development. Neurobiol Dis, 2019. 122: p. 49–63.

52. Fulmer, C.G., et al., Astrocyte-derived BDNF supports myelin protein synthesis after cuprizone-induced demyelination. J Neurosci, 2014. 34(24): p. 8186–96.

53. Clarke, L.E., et al., Normal aging induces A1-like astrocyte reactivity. Proc Natl Acad Sci U S A, 2018. 115(8): p. E1896–E1905.

54. Kostuk, E.W., J. Cai, and L. Iacovitti, Subregional differences in astrocytes underlie selective neurodegeneration or protection in Parkinson’s disease models in culture. Glia, 2019. 67(8): p. 1542–1557.

55. O’Malley, E.K., et al., Mesencephalic type I astrocytes mediate the survival of substantia nigra dopaminergic neurons in culture. Brain Res, 1992. 582(1): p. 65–70.

56. Di Malta, C., et al., Astrocyte dysfunction triggers neurodegeneration in a lysosomal storage disorder. Proc Natl Acad Sci U S A, 2012. 109(35): p. E2334–42.

57. Moruno-Manchon, J.F., et al., Sphingosine kinase 1-associated autophagy differs between neurons and astrocytes. Cell Death Dis, 2018. 9(5): p. 521.

58. Lieberman, O.J., et al., Cell-type-specific regulation of neuronal intrinsic excitability by macroautophagy. Elife, 2020. 9.

59. Vijayan, V. and P. Verstreken, Autophagy in the presynaptic compartment in health and disease. J Cell Biol, 2017. 216(7): p. 1895–1906.

60. Lieberman, O.J. and D. Sulzer, The Synaptic Autophagy Cycle. J Mol Biol, 2020. 432(8): p. 2589–2604.

61. Schildge, S., et al., Isolation and culture of mouse cortical astrocytes. J Vis Exp, 2013(71).

62. Pan, P.Y. and T.A. Ryan, Calbindin controls release probability in ventral tegmental area dopamine neurons. Nat Neurosci, 2012. 15(6): p. 813–5.

